# Growth arrest of *Staphylococcus aureus* induces daptomycin tolerance via cell wall remodelling

**DOI:** 10.1101/2022.08.10.503462

**Authors:** Elizabeth V. K. Ledger, Andrew M. Edwards

## Abstract

Almost all bactericidal drugs require bacterial replication and/or metabolic activity for their killing activity. When these processes are inhibited by bacteriostatic antibiotics, bacterial killing is significantly reduced. One notable exception is the lipopeptide antibiotic daptomycin, which has been reported to efficiently kill non-dividing bacteria. However, these studies employed only brief periods of growth arrest. We found that a bacteriostatic concentration of the protein synthesis inhibitor tetracycline led to a time-dependent induction of daptomycin tolerance in *S. aureus*, with ^~^100,000-fold increase in survival after 16 h growth arrest relative to exponential phase bacteria. Daptomycin tolerance required glucose and was associated with increased production of the cell wall polymers peptidoglycan and wall-teichoic acids. However, whilst accumulation of peptidoglycan was required for daptomycin tolerance, only a low abundance of wall teichoic acid was necessary. Therefore, whilst tolerance to most antibiotics occurs passively due to a lack of metabolic activity and/or replication, daptomycin tolerance arises via active cell wall remodelling.

## Introduction

All antibiotics disrupt essential processes or structures in bacteria [1,2]. Whether this disruption leads to bacterial killing or growth inhibition is a function of antibiotic class, as well as the physiological state of the cell [1,2,3,4,5,6,7,8,9,10,11]. Understanding the factors that modulate antibiotic activity is important because the inability of bactericidal antibiotics to kill bacteria has been linked to treatment failure and the emergence of antibiotic resistance [12,13,14,15,16,17]. This is particularly important in the case of *S. aureus*, since this pathogen causes several refractory infections, including bacteraemia, infective endocarditis, osteomyelitis and implant infections [12,13,14,18].

It is well established that growth arrest of bacteria, for example via exposure to a bacteriostatic antibiotic, significantly increases bacterial survival during subsequent exposure to most bactericidal antibiotics [3,8,9,10]. However, the reasons for this are subject to debate because, in addition to blocking replication, bacteriostatic antibiotics also inhibit bacterial respiration, which in turn compromises metabolic pathways [10]. Since a lack of metabolic activity results in growth arrest it has been challenging to determine whether killing by bactericidal antibiotics requires active metabolism or replication (or both) to occur. However, a recent study provided evidence that metabolic activity is more important than replication for killing of the human and animal pathogen *Staphylococcus aureus* by several different antibiotics, and an earlier report linked a lack of cellular ATP in this bacterium to antibiotic tolerance [8,11]. Since the physiological state of bacteria at infection sites is likely to be different from that in laboratory medium, understanding the host and bacterial factors that modulate antibiotic susceptibility may lead to better treatment outcomes [12,13].

One of the very few antibiotics reported to efficiently kill growth-arrested *S. aureus* is the lipopeptide daptomycin, which is used to treat infections caused by drug-resistant Gram-positive pathogens such as Methicillin Resistant *Staphylococcus aureus* (MRSA) [19,20,21]. Daptomycin targets phosphatidylglycerol (PG) and lipid II in the membrane, where it forms oligomeric complexes leading to membrane depolarisation and permeabilization [19,22,23,24]. The binding of daptomycin to lipid II also disrupts cell wall synthesis, which is further compromised by mis-localisation of enzymes involved in peptidoglycan synthesis [19,24,25,26].

The ability of daptomycin to kill growth-arrested bacteria has been demonstrated using cells pre-treated with bacteriostatic antibiotics, membrane depolarising agents or cold temperature, or those in stationary phase, conditions that almost completely block killing by other bactericidal antibiotics [10,20,21]. However, whilst daptomycin maintains bactericidal activity against non-replicating bacteria, these and other studies clearly showed that the degree and rate of killing of growth-arrested bacteria was reduced compared to replicating bacteria [10,20,21,27,28,29].

The reason for the reduced susceptibility of non-replicating *S. aureus* to killing by daptomycin is unknown [11] but may have important clinical implications because the potent activity of this antibiotic *in vitro* is not always replicated *in vivo*. In a retrospective analysis, >25% patients with staphylococcal bacteraemia given daptomycin monotherapy died [30], whilst other studies have reported treatment failure rates of 11-25% [31,32,33], demonstrating that there is a pressing need to improve daptomycin-based treatment approaches.

Therefore, the aims of this work were to determine the degree to which growth arrest confers daptomycin tolerance, the mechanism responsible and to identify ways to prevent tolerance and enhance daptomycin treatment efficacy.

## Results

### Antibiotic-mediated growth arrest induces daptomycin tolerance in a time- and nutrient-dependent manner

To test whether growth arrest of *S. aureus* led to daptomycin tolerance we grew bacteria to mid-exponential phase, and then exposed them directly to daptomycin, or incubated cells with a bacteriostatic concentration of tetracycline for 16 h before exposure to daptomycin. As expected, exponential phase *S. aureus* was very susceptible to daptomycin, with a ~6 log reduction in CFU counts after 6 h antibiotic exposure (Fig. 1A). By contrast, growth arrest with tetracycline completely prevented daptomycin killing (Fig. 1A).

**Figure 1.**
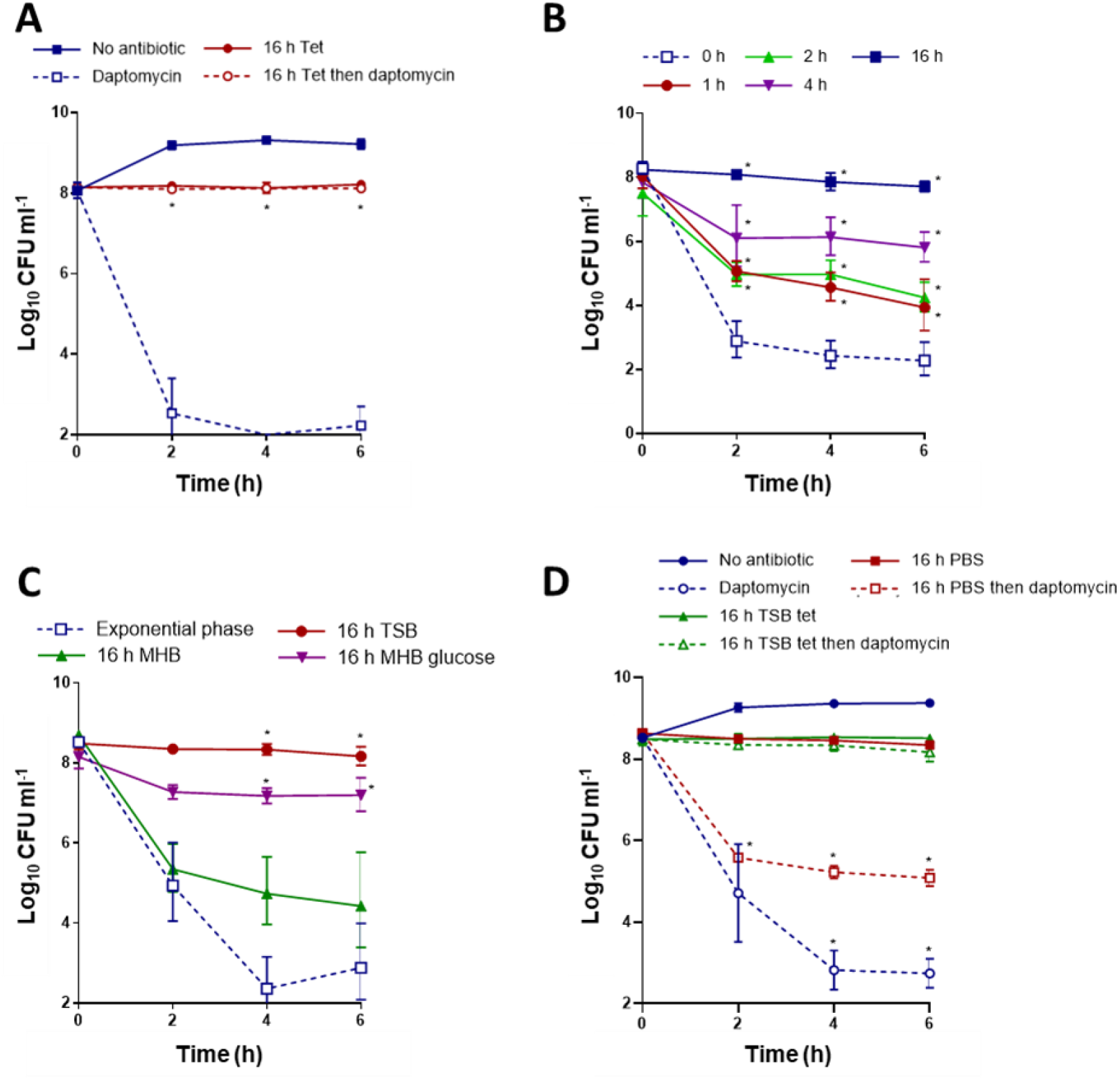
Tetracycline-mediated growth arrest induces daptomycin tolerance in a time- and nutrient-dependent manner. (**A**) *S. aureus* USA300 LAC* was grown to mid-exponential phase, or grown to mid-exponential phase and then incubated for 16 h with 1.25 μg ml^-1^ tetracycline, and then log_10_ CFU ml^-1^ determined throughout a 6 h exposure to 20 μg ml^-1^ daptomycin. (**B**) *S. aureus* USA300 LAC* was grown to mid-exponential phase and incubated for indicated lengths of time with 1.25 μg ml^-1^ tetracycline before log_10_ CFU ml^-1^ were determined throughout a 6 h exposure to 20 μg ml^-1^ daptomycin. (**C**) *S. aureus* USA300 LAC* was grown to mid-exponential phase or incubated for 16 h with 1.25 μg ml^-1^ tetracycline in TSB, MHB or MHB supplemented with 2.5 g L^-1^ glucose, before being exposed to 20 μg ml^-1^ daptomycin in TSB for 6 h and log_10_ CFU ml^-1^ determined. (**D**) Log_10_ CFU ml^-1^ of mid-exponential phase cells, or cultures incubated in TSB supplemented with 1.25 μg ml^-1^ tetracycline or in PBS for 16 h, during a 6 h exposure to 20 μg ml^-1^ daptomycin in TSB. Data represent the geometric mean ± geometric standard deviation of three independent experiments. Data in **A** were analysed by two-way ANOVA with Sidak’s *post-hoc* test (* P < 0.05; daptomycin-treated tetracycline-arrested vs exponential phase). Data in **B** were analysed by two-way ANOVA with Tukey’s *post-hoc* test (* P < 0.05; non-arrested vs tetracycline-arrested). Data in **C** were analysed by two-way ANOVA with Dunnett’s *post-hoc* test (* P < 0.05; exponential phase vs tetracycline-arrested). Data in **D** were analysed by two-way ANOVA with Dunnett’s *post-hoc* test (* P < 0.05; tetracycline-arrested vs non-arrested/PBS-arrested. Tet, tetracycline.

Since the 16 h of growth arrest used here was longer than that used in previous studies [10,20,21], we next examined how the duration of growth inhibition affected daptomycin tolerance. In keeping with previous work [10,20,21], a short period of growth inhibition (1 or 2 h) reduced daptomycin killing by ~100-fold relative to growing bacteria (Fig. 1B). Cells that were growth-arrested for 4 h showed a further reduction in susceptibility to daptomycin, with 4 log higher survival relative to growing *S. aureus* cells, whilst a 16 h incubation in tetracycline again conferred complete protection from daptomycin-mediated killing (Fig. 1B).

Since the induction of daptomycin tolerance was time-dependent, we hypothesised that an active process was required for the loss of antibiotic susceptibility. Therefore, we investigated the nutritional requirements for tolerance by arresting growth for 16 h with tetracycline in either tryptic soy broth (TSB) or Müller-Hinton broth (MHB), media with different nutrient compositions.

Whilst 16 h growth arrest in TSB completely blocked killing by daptomycin, this was not the case with MHB, with >1000-fold reduction in CFU after 6 h exposure to the lipopeptide antibiotic (Fig. 1C). One of the differences between TSB and MHB is that the former contains glucose (2.5 g L^-1^), and supplementation of MHB with this sugar demonstrated that it was crucial for the induction of tolerance (Fig. 1C). To confirm the requirement for nutrients, and to demonstrate daptomycin tolerance was not simply due to a lack of growth or metabolic activity, we arrested growth via nutrient limitation by incubating bacteria for 16 h in phosphate-buffered saline (PBS) (Fig. 1D). Nutrient limitation failed to provide the high level of daptomycin tolerance seen in TSB, with a 3 log reduction of CFU counts within 2 h of daptomycin exposure (Fig. 1D).

Together, these data show that growth arrest leads to very high levels of daptomycin tolerance via a time- and nutrient-dependent process, in contrast to the tolerance of other bactericidal antibiotics, which is linked to a lack of metabolic activity [7,9,11].

### Daptomycin tolerance requires synthesis of cell surface components

Next, we aimed to determine which biosynthetic pathways were required for daptomycin tolerance by investigating the effects of growth arrest by various classes of antibiotic. To do this, we incubated exponential phase cultures of *S. aureus* for 16 h with growth inhibitory concentrations of chloramphenicol (inhibitor of protein synthesis), ciprofloxacin (inhibitor of DNA replication), cephalexin (inhibitor of cell wall synthesis), tunicamycin (inhibitor of wall teichoic acid (WTA) synthesis) or AFN-1252 (inhibitor of fatty acid synthesis), and then challenged with daptomycin.

Growth arrest with each of the antibiotics led to varying degrees of daptomycin tolerance (Fig. 2A – E). Similarly to tetracycline, inhibition of protein synthesis with chloramphenicol led to very high levels of tolerance, with daptomycin killing less than 1 log of *S. aureus* CFU (Fig. 2A). Growth arrest with ciprofloxacin also led to a high degree of tolerance (Fig. 2B). However, growth arrest with antibiotics that inhibited synthesis of components of the cell surface (peptidoglycan, WTA or fatty acids) resulted in lower levels of daptomycin tolerance (Fig. 2C, D, E). This was especially pronounced with cephalexin and tunicamycin, where daptomycin caused 2-3 log reductions on CFU within 6 h (Fig. 2C, D).

**Figure 2.**
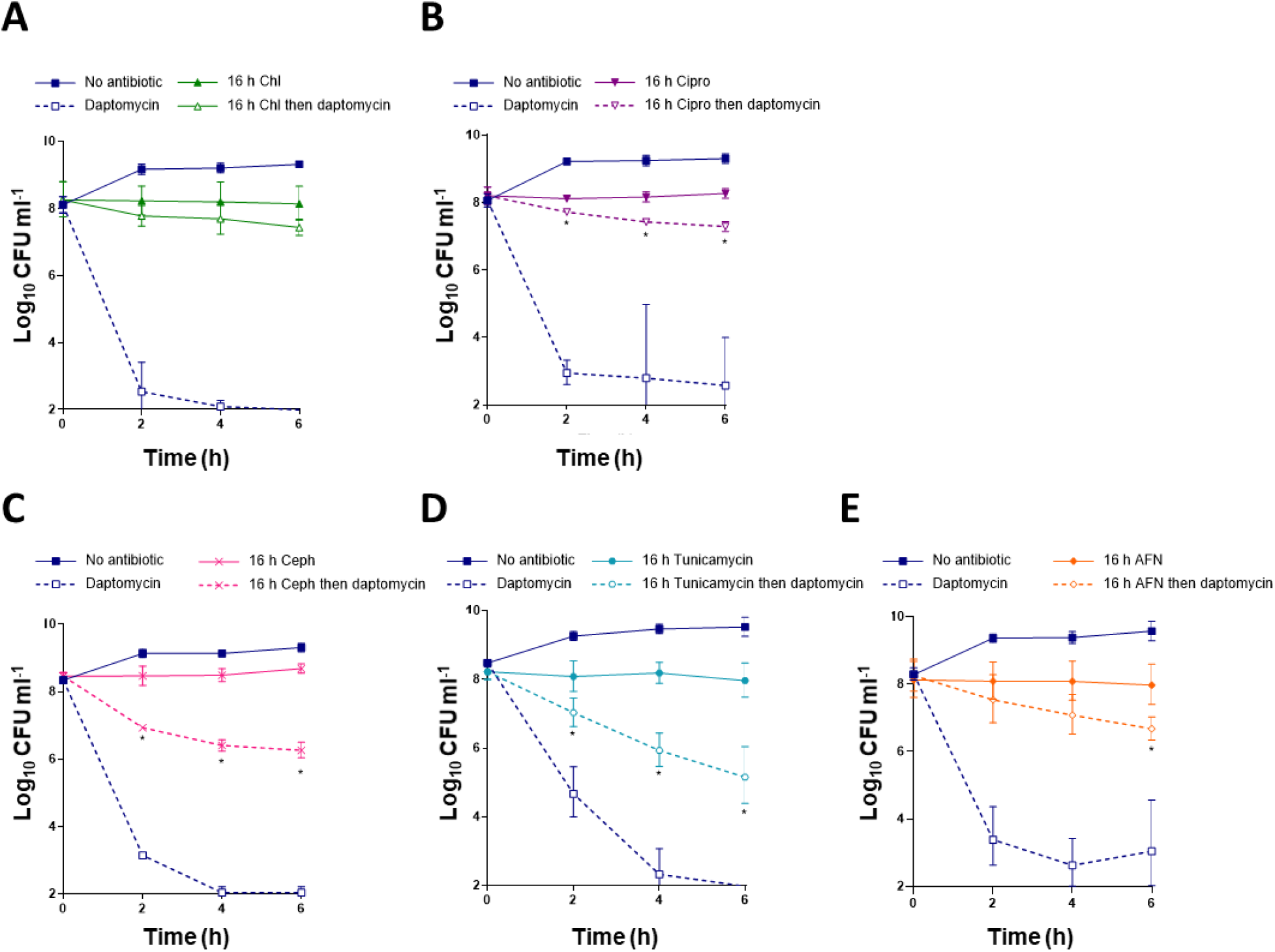
Antibiotics with different mechanisms of action induce tolerance to different degrees. *S. aureus* USA300 LAC* were grown to mid-exponential phase or incubated for 16 h with (**A**) 20 μg ml^-1^ chloramphenicol, (**B**) 160 μg ml^-1^ ciprofloxacin, (**C**) 32 μg ml^-1^ cephalexin, (**D**) 0.5 μg ml^-1^ tunicamycin or (**E**) 0.15 μg ml^-1^ AFN-1252 before log_10_ CFU ml^-1^ were determined throughout a 6 h exposure to 20 μg ml^-1^ daptomycin. Data represent the geometric mean ± geometric standard deviation of three independent experiments. Data were analysed by two-way ANOVA with Sidak’s *post-hoc* test (* P < 0.05; daptomycin-treated antibiotic-arrested vs daptomycin-treated exponential phase). Chl, chloramphenicol; Cipro, ciprofloxacin; Ceph, cephalexin; AFN, AFN-1252.

Combined, these data show that growth arrest by different classes of antibiotics resulted in varying degrees of daptomycin tolerance, but full tolerance required peptidoglycan, WTA and fatty acids, suggesting that changes to the bacterial cell surface contributed to tolerance.

### Antibiotic-mediated growth arrest reduces daptomycin binding and membrane damage

To understand how growth arrest-induced changes to the cell wall reduced daptomycin susceptibility, we next tested whether antibiotic-mediated growth arrest affected the ability of daptomycin to bind to its membrane target.

To do this, we measured attachment of fluorescent BoDipy-daptomycin to mid-exponential phase or tetracycline-arrested cultures over a 6 h period, with aliquots taken every 2 h to determine the kinetics of antibiotic binding. In line with the bacterial survival data, exponential phase cells were bound by high levels of daptomycin within 2 h, whereas growth-arrested cells had significantly reduced levels of bound daptomycin (Fig. 3A).

**Figure 3.**
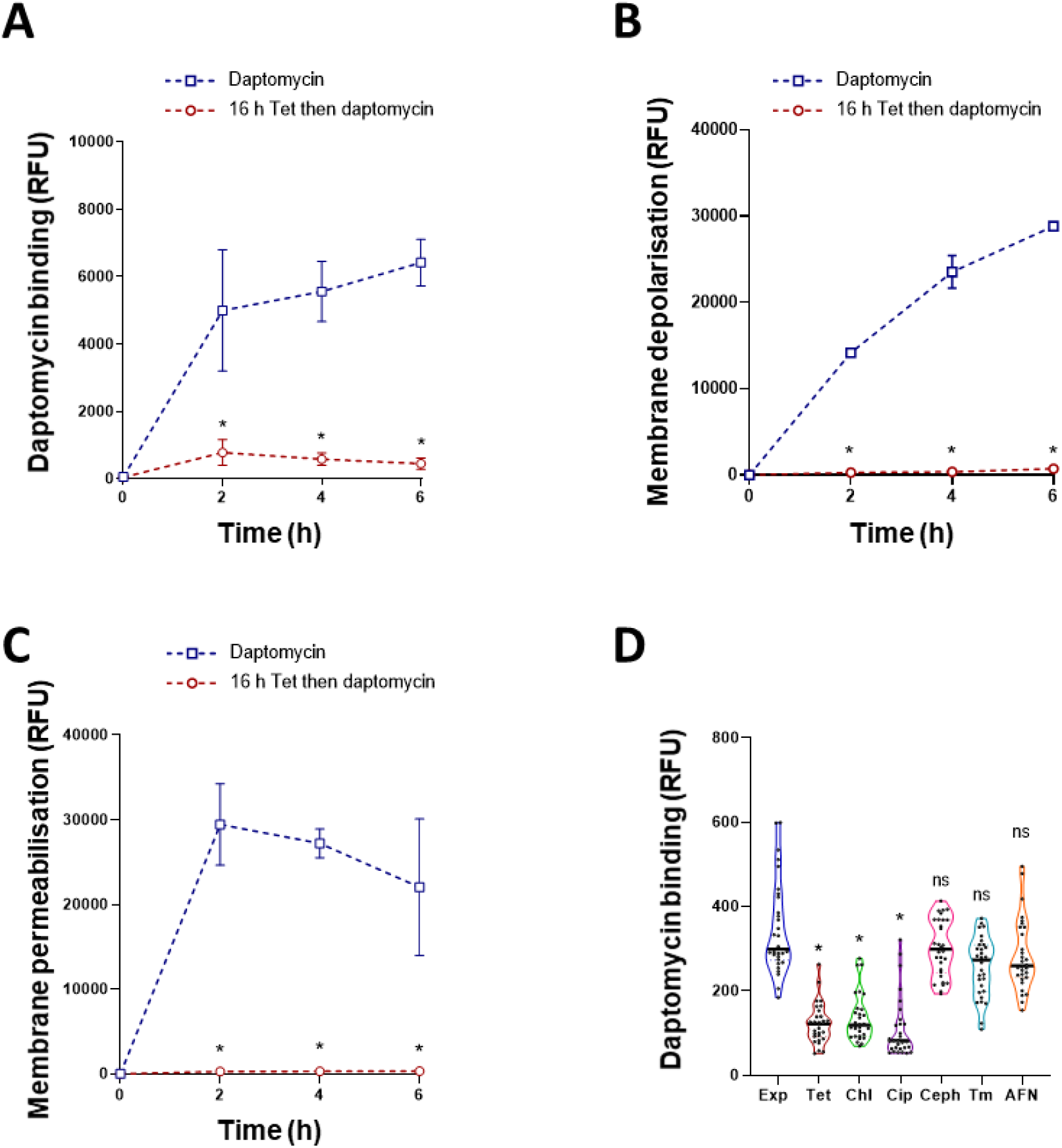
Antibiotic-mediated growth arrest reduces daptomycin binding and membrane damage. (**A**) *S. aureus* USA300 LAC* were grown to mid-exponential phase or incubated for 16 h with 1.25 μg ml^-1^ tetracycline before being exposed to 80 μg ml^-1^ BoDipy-daptomycin and cell-associated fluorescence determined. (**B**) DiSC_3_(5) and (**C**) propidium iodide fluorescence of mid-exponential or tetracycline-arrested cultures during a 6 h exposure to 20 μg ml^-1^ daptomycin. (**D**) BoDipy-daptomycin binding to exponential phase and antibiotic-arrested cultures of *S. aureus* USA300 LAC*. Cultures were incubated with 80 μg ml^-1^ BoDipy-daptomycin for indicated lengths of time, washed, fixed and analysed by fluorescence microscopy. The fluorescence of 30 cells per condition was measured. Data in **A** – **C** represent the mean ± standard deviation of three independent experiments. Data in **D** represent individual cellular measurements with the median indicated. Data in **A** – **C** were analysed by two-way ANOVA with Tukey’s *post-hoc* test (* P < 0.05; tetracycline-arrested vs exponential phase). Data in **D** were analysed by Kruskal-Wallis test with Dunn’s *post-hoc* test (* P < 0.05; Exp vs antibiotic-arrested). Exp, exponential phase; Tet, tetracycline; Chl, chloramphenicol; Cip, ciprofloxacin; Ceph, cephalexin; Tm, tunicamycin; AFN, AFN-1252.

We next measured membrane depolarisation and permeability using the fluorescent dyes DiSC3(5) and propidium iodide, respectively. As expected from previous work [34], the membranes of exponential phase *S. aureus* were rapidly depolarised and permeabilised on exposure to daptomycin (Fig. 3B, C). By contrast, no membrane disruption was detected by either assay in the tetracycline-arrested cultures (Fig. 3B, C).

Finally, we investigated the effect of growth arrest by other classes of antibiotic on daptomycin binding. As described above, growth arrest with tetracycline significantly reduced daptomycin binding (Fig. 3D). In agreement with the data from bacterial survival assays (Fig. 2A, B, C, D, E), chloramphenicol and ciprofloxacin also significantly reduced daptomycin binding compared to exponential phase cultures, while BoDipy-daptomycin bound as well to cells incubated with cephalexin, tunicamycin or AFN-1252 as to exponential phase cultures (Fig. 3D).

Taken together, these data demonstrate that the daptomycin tolerance of growth-arrested bacteria is due to reduced binding of the lipopeptide to its membrane targets, likely resulting from changes to the cell envelope.

### Daptomycin tolerance requires changes to the cell wall but not the membrane

The next objective was to determine which components of the cell surface were required for daptomycin tolerance. As changes to both the cell wall and membrane components of *S. aureus* have previously been associated with reduced daptomycin susceptibility [19,35,36,37,38,39,40,41,42], we tested each in turn, starting with membrane phospholipid synthesis.

The main phospholipid species in the *S. aureus* membrane are phosphatidylglycerol (PG), lysyl-phosphatidylglycerol (LPG) and cardiolipin (CL) [37,39]. As PG is essential, we investigated whether synthesis of either LPG (by MprF) or CL (by either Cls1 or Cls2) were required for tolerance using mutants defective for these proteins from the NARSA library, which was generated in the USA300 LAC JE2 background. Therefore, we first confirmed that tetracycline induced daptomycin tolerance in the JE2 WT strain (Fig. 4A), before testing the phenotypes of each of the mutants. Similar to the WT strain, growth arrest conferred daptomycin tolerance in each of the three phospholipid synthase mutants, with no significant killing observed in any of the strains (Fig. 4B, C, D).

**Figure 4.**
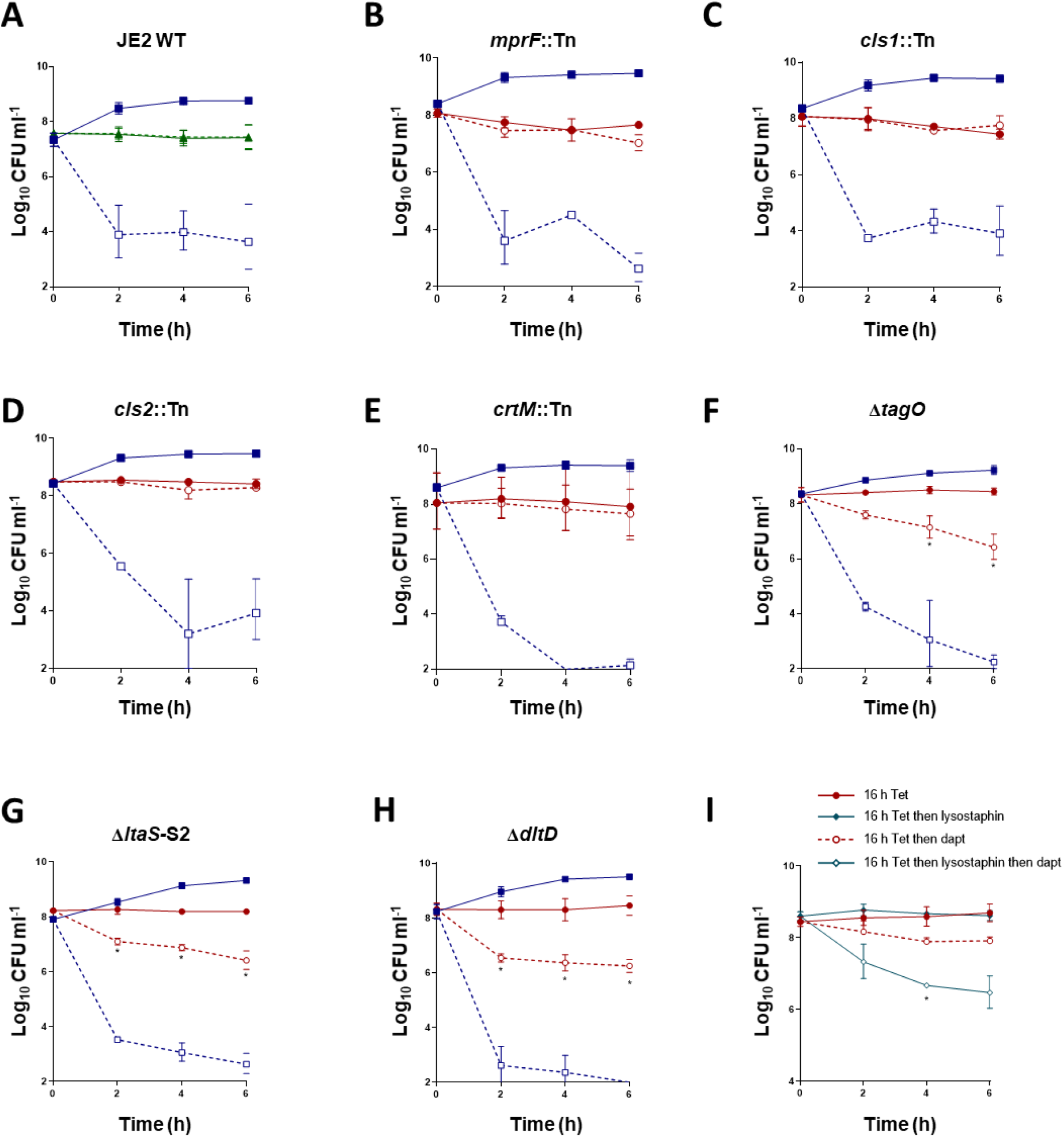
Daptomycin tolerance requires changes in the cell wall and not the cell membrane. Log_10_ CFU ml^-1^ of exponential phase and tetracycline-arrested cultures of (**A**) JE2 WT, (**B**) JE2 *mprF*::Tn, (**C**) JE2 *cls1::Tn*, (**D**) JE2 *cls2*::Tn, (**E**) JE2 *crtM*::Tn, (**F**) LAC* Δ*tagO*, (**G**) LAC* Δ*ltaS-S2*, (**H**) LAC* Δ*dltD* during a 6 h exposure to 20 μg ml^-1^ daptomycin. (**I**) Log_10_ CFU ml^-1^ of tetracycline-arrested cultures of *S. aureus* LAC*, after partial digestion of the cell wall with lysostaphin, or not, and during a 6 h exposure to 20 μg ml^-1^ daptomycin. Data in represent the geometric mean ± geometric standard deviation of three independent experiments. Data in **A** – **H** were analysed by two-way ANOVA with Sidak’s *post-hoc* test (* P < 0.05; tetracycline-arrested vs tetracycline-arrested + daptomycin). Data in **I** were analysed by two-way ANOVA with Dunnett’s *post-hoc* test (* P < 0.05; lysostaphin treated vs untreated). Tet, tetracycline, Dapt, daptomycin.

During our assays, we observed that growth-arrested *S. aureus* were much more strongly pigmented than exponential phase cultures. This pigment, staphyloxanthin, is a major factor influencing membrane fluidity and has been implicated in reducing daptomycin susceptibility [42]. Therefore, we next tested a mutant defective in pigment synthesis, *crtM::Tn*. However, despite lacking staphyloxanthin, this mutant also showed high levels of daptomycin tolerance after being incubated with tetracycline, confirming that synthesis of this pigment did not contribute to bacterial survival (Fig. 4E). Therefore, we found no evidence that daptomycin tolerance was due to changes in membrane phospholipid composition or staphyloxanthin content.

A major component of the cell envelope are teichoic acids, which can either be covalently bound to the cell wall (WTA) or anchored to the membrane (lipoteichoic acid, LTA) [43]. These polymers are both modified with D-alanine groups by the products of the *dltABCD* operon, reducing the net negative charge of the surface [43]. To test whether either of these polymers were required for tolerance we examined mutants lacking WTA or LTA (Δ*tagO* or Δ*ltaS*-S2, respectively) or lacking the D-alanine modification (Δ*dltD*). These assays showed that survival of each mutant during daptomycin exposure was ~2 log lower than that of the WT after 6 h, indicating that the presence of D-alanylated teichoic acids was required for full tolerance (Fig. 4F, G, H).

To further test whether tolerance required the cell wall, we arrested growth of *S. aureus* with tetracycline to induce tolerance, then partially digested peptidoglycan with lysostaphin and measured daptomycin susceptibility. Lysostaphin treatment led to a ~2 log reduction in bacterial survival compared to cells with an intact cell wall, confirming a key role for the cell wall in mediating daptomycin tolerance induced by growth arrest (Fig. 4I).

Taken together, the induction of daptomycin tolerance in growth-arrested cells was due to changes to the cell wall but not the membrane.

### Growth arrest causes peptidoglycan and WTA accumulation

Having established that the cell wall was required for growth-arrest induced daptomycin tolerance, we next aimed to determine the nature of the changes associated with tolerance.

Firstly, we looked at whether growth arrest led to increases in any of the three main components of the wall, peptidoglycan, WTA and LTA. To test whether peptidoglycan was accumulating, we used HADA, a fluorescent D-amino acid that is incorporated into peptidoglycan as it is synthesised [44]. Growth arrest was carried out in TSB supplemented with HADA for various lengths of time before bacteria were imaged and the fluorescence quantified by microscopy. This demonstrated that longer periods of growth arrest were associated with higher levels of HADA fluorescence, indicating that peptidoglycan was accumulating over time (Fig. 5A). Next, we extracted and quantified WTA from exponential phase and growth-arrested cultures, demonstrating a time-dependent increase in the quantity of WTA, with three times more WTA present after a 16 h growth arrest than in exponential phase (Fig. 5B). By contrast, extraction and detection of LTA by western blotting revealed no significant increase in the quantity of LTA in growth-arrested cells compared to those in exponential phase (Fig. 5C).

**Figure 5.**
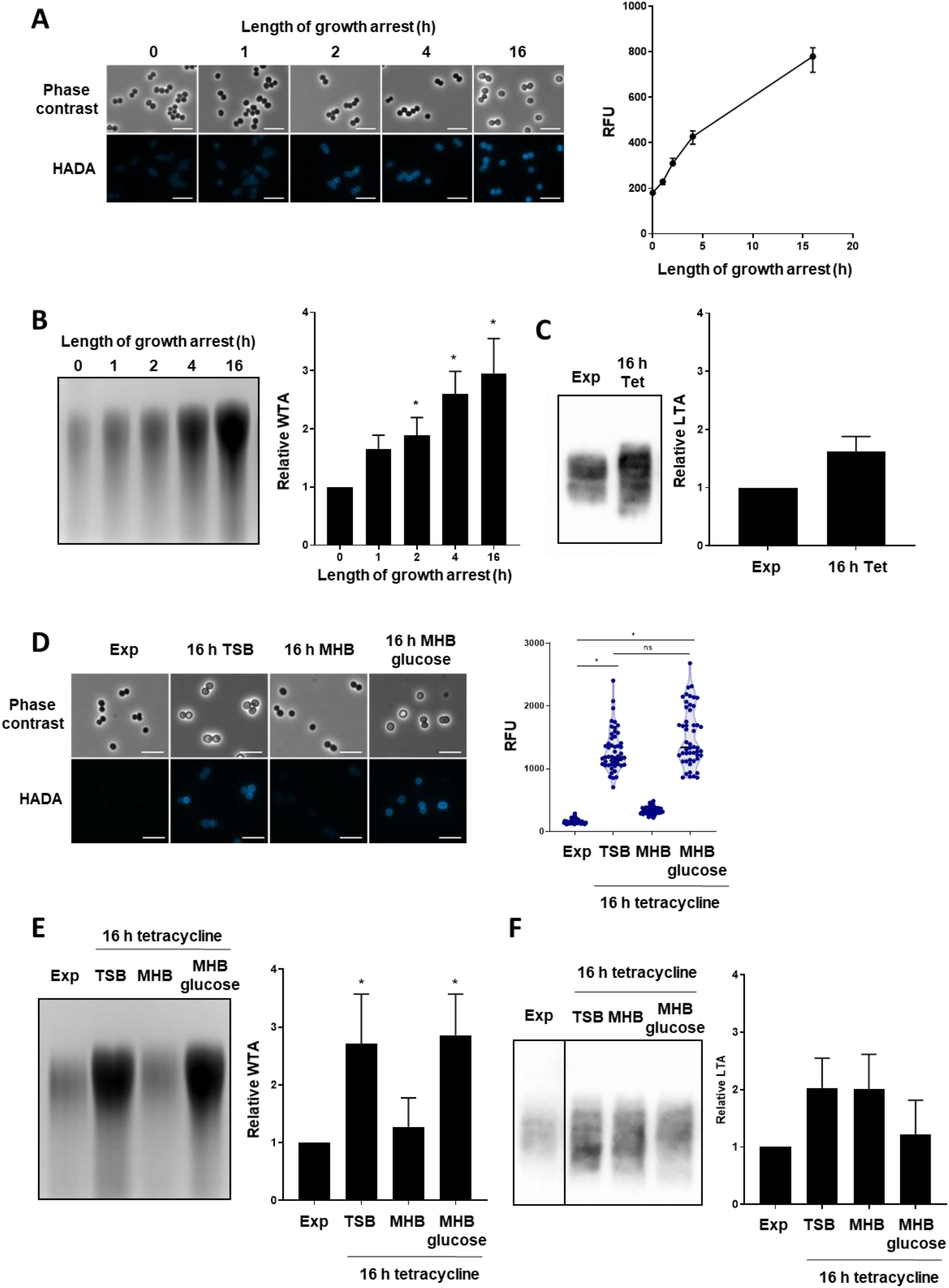
Growth arrest with tetracycline leads to increases in peptidoglycan and WTA. (**A**) Exponential phase *S. aureus* LAC* were growth-arrested with tetracycline for indicated lengths of time in the presence of 25 μM HADA before being analysed by phase contrast and fluorescence microscopy and the fluorescence of the cell surface of individual cells quantified. Exponential phase *S. aureus* LAC* were growth-arrested with tetracycline for indicated lengths of time before (**B**) WTA was extracted, analysed by native PAGE, visualised with alcian blue and quantified or (**C**) LTA was analysed by western blotting and quantified. Data are expressed as relative to exponential phase values. (**D**) Tetracycline-mediated growth arrest was carried out in TSB, MHB or MHB supplemented with 2.5 g L^-1^ glucose for 16 h in the presence of 25 μM HADA before cells were visualised by phase contrast and fluorescence microscopy and the fluorescence of the surface of individual cells quantified. Tetracycline-mediated growth arrest was carried out in TSB, MHB or MHB supplemented with 2.5 g L^-1^ glucose for 16 h before (**E**) WTA was extracted, analysed by PAGE and quantified with alcian blue staining and (**F**) LTA was extracted, analysed by western blotting and quantified. Data are expressed as relative to exponential phase values. Data in **B**, **C**, **E** and **F** represent the mean ± standard deviation of three independent experiments. Data in **A** represent the median fluorescence ± 95 % confidence intervals of 50 individual cells. Data in **D** represent the fluorescence values of individual cells with the median indicated. WTA PAGE and LTA western blots were performed three times and representative images are shown. Data in **B** were analysed by one-way ANOVA with Tukey’s *post-hoc* test (* P < 0.05; tetracycline-arrested vs exponential phase). Data in **C** were analysed by t-test (no significant difference). Data in **D** were analysed by Kruskal-Wallis with Dunn’s *post-hoc* test (* P < 0.05). Data in **E** and **F** were analysed by two-way ANOVA with Dunnett’s *post-hoc* test (* P < 0.05; tetracycline-arrested vs exponential). No significant differences were observed in panel **F**. Exp, exponential phase; Tet, tetracycline; TSB, tryptic soy broth; MHB, müller-hinton broth.

Having identified that growth arrest led to time-dependent increases in both peptidoglycan and WTA, but not LTA, we next investigated whether the increases in these polymers were nutrient-dependent. In line with the tolerance data (Fig. 1B), both peptidoglycan and WTA abundance increased when growth arrest occurred in TSB, whereas no significant accumulation of either polymer was observed after growth arrest in MHB (Fig. 5D, E). Furthermore, growth arrest in MHB supplemented with glucose led to increased levels of both peptidoglycan and WTA (Fig. 5D, E). As confirmation that increased LTA was not associated with growth arrest-induced daptomycin tolerance, we found no differences between the levels of LTA in cells arrested in TSB or MHB, and supplementation of MHB with glucose did not lead to increased LTA (Fig. 5F).

Taken together, tetracycline-mediated growth arrest leads to time- and nutrient-dependent increases in peptidoglycan and WTA, but not LTA.

### Daptomycin tolerance is due to the accumulation of peptidoglycan but not WTA

The final objective was to determine whether the accumulation of either peptidoglycan or WTA that occurred during growth arrest was responsible for tolerance.

To do this, we growth-arrested bacteria with tetracycline in TSB in the presence of inhibitors of WTA synthesis, teichoic acid D-alanylation or peptidoglycan synthesis and then measured daptomycin tolerance. Firstly, showed that tunicamycin blocked the increase in WTA that occurred during growth arrest with tetracycline, but maintained levels seen in exponential cells (Fig. 6A). We then measured the daptomycin susceptibility of the tetracycline plus tunicamycin-arrested bacteria. Despite having the same levels of WTA as exponential phase bacteria, these cells had very high levels of daptomycin tolerance, with no significant killing by 6 h. Therefore, the accumulation of WTA that occurs during growth arrest was not required for daptomycin tolerance (Fig. 6B).

**Figure 6.**
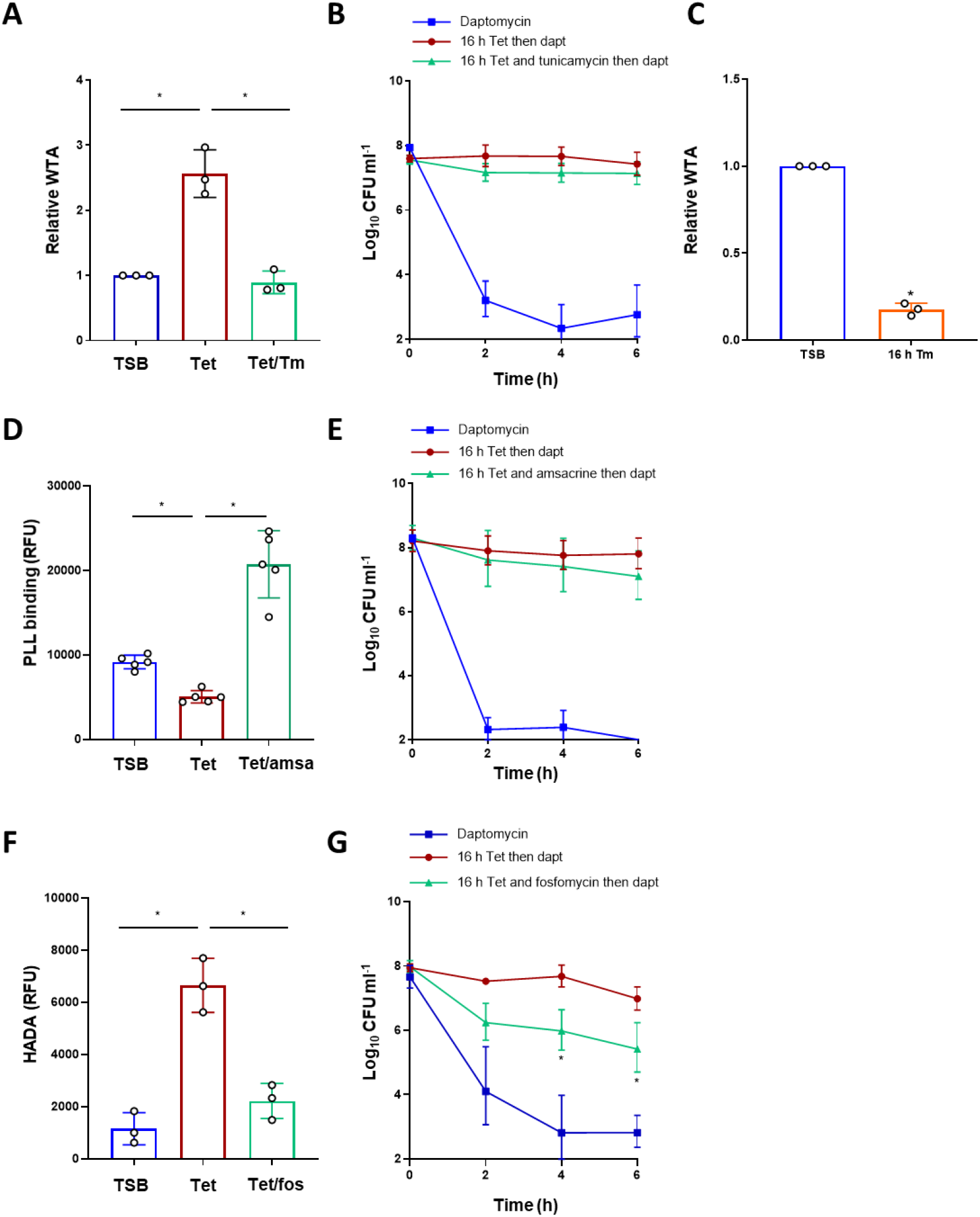
Daptomycin tolerance is due to the accumulation of peptidoglycan but not WTA. Exponential phase *S. aureus* LAC* were growth-arrested with tetracycline +/− 0.5 μg ml^-1^ tunicamycin for 16 h before (**A**) WTA was extracted, analysed by PAGE and quantified by alcian blue staining or (**B**) log_10_ CFU ml^-1^ determined during a 6 h exposure to 20 μg ml^-1^ daptomycin. Exponential phase *S. aureus* were growth-arrested with 0.5 μg ml^-1^ tunicamycin for 16 h, or not, before WTA was extracted, analysed by PAGE and quantified by alcian blue staining (**C**). Exponential phase *S. aureus* LAC* were growth-arrested with tetracycline +/− 10 μg ml^-1^ amsacrine for 16 h before (**D**) surface charge was measured using FITC-PLL or (**E**) log_10_ CFU ml^-1^ determined during a 6 h exposure to 20 μg ml^-1^ daptomycin. Exponential phase *S. aureus* LAC* were growth-arrested with tetracycline +/− 8 μg ml^-1^ fosfomycin for 16 h before (**F**) peptidoglycan content was measured using HADA or (**G**) log_10_ CFU ml^-1^ determined during a 6 h exposure to 20 μg ml^-1^ daptomycin. Data in **A**, **C**, **D** and **F** represent the mean ± standard deviation of indicated number of independent experiments. Data in **B**, **E** and **G** represent the geometric mean ± geometric standard deviation of three independent experiments. Data in **A**, **D** and **F** were analysed by one-way ANOVA with Tukey’s *post-hoc* test (* P < 0.05). Data in **B**, **E** and **G** were analysed by two-way ANOVA with Dunnett’s *post-hoc* test (* P < 0.05; tetracycline-arrested vs tetracycline + inhibitor-arrested). Data in **C** were analysed by Student’s t-test (* P < 0.05) TSB, tryptic soy broth; Tet, tetracycline; Tm, tunicamycin; Amsa, amsacrine; Fos, fosfomycin; Dapt, daptomycin.

Since we’d previously found that cells growth arrested with tunicamycin alone had a low level of tolerance (Fig. 2D) we examined the levels of WTA in these bacteria. This showed that cells that were growth arrested with tunicamycin alone had ^~^5-fold lower levels of WTA relative to exponential-phase cells (Fig. 6C) or cells growth arrested with tetracycline plus tunicamycin (Fig. 6A). These data, together with those obtained with the WTA-deficient Δ*tagO* mutant (Fig. 4F), led us to conclude that a minimal amount of WTA is needed for tolerance, but that accumulation of the polymer above levels seen in exponential-phase cells was not required.

Next, we assessed the importance of D-alanylation of teichoic acids for daptomycin tolerance using amsacrine, which inhibits DltB [45]. We showed that this inhibitor worked under the conditions used by measuring the binding of fluorescently labelled poly-L-lysine (FITC-PLL), a positively charged polymer which is an indicator of bacterial surface charge (Fig. 6D). Exposure of bacteria treated with tetracycline and amsacrine to daptomycin revealed a very high level of tolerance, demonstrating that the increased levels of teichoic acid D-alanylation seen in growth arrested cells did not contribute to antibiotic survival (Fig. 6E), although the presence of some D-alanylation is required (Fig. 4H)

Finally, we investigated whether the growth arrest-induced accumulation of peptidoglycan mediated daptomycin tolerance. Peptidoglycan accumulation was inhibited using fosfomycin, which prevented the increase in HADA incorporation seen during tetracycline-mediated growth arrest (Fig. 6F). In contrast to tetracycline-arrested cultures, bacteria where peptidoglycan accumulation had been prevented were significantly killed by daptomycin, with over 2 log reduction in CFU counts after by 6 h exposure to the lipopeptide antibiotic (Fig. 6G).

Therefore, while the presence of both WTA and its D-alanine modification are required for daptomycin tolerance (Fig. 4F, G, H), it is the accumulation of peptidoglycan, and not WTA, which is responsible for tolerance.

## Discussion

In contrast to most bactericidal antibiotics, daptomycin has been reported to kill growth-arrested bacteria, including those exposed to bacteriostatic drugs [10,20,21]. However, previous work has shown that brief periods of antibiotic-mediated growth arrest reduced daptomycin-mediated killing, suggesting that the bactericidal activity of the lipopeptide antibiotic is affected by growth status. The aims of this work were to understand the degree to which growth arrest can reduce killing by daptomycin and the mechanism responsible.

Several studies have shown that antibiotic tolerance arises due to a lack of metabolic activity and/or replication reducing the deleterious effects of bactericidal antibiotics such as the production of reactive oxygen species [7,8,9,10,11,15]. By contrast, in this work, we show that growth arrest induces very high levels of daptomycin tolerance via an active mechanism that requires time, nutrients and peptidoglycan synthesis.

Tetracycline-mediated growth arrest led to an accumulation of major cell wall components, which agrees with previous work showing that inhibition of protein synthesis halts growth but not peptidoglycan biosynthesis in *S. aureus* [46]. However, whilst the presence of WTA and D-alanylation were essential for tolerance, their accumulation was not. This is in keeping with previous work from our group, that showed that peptidoglycan accumulation in bacteria incubated in serum was dependent upon the presence, but not accumulation, of D-alanylated WTA [34]. Increased levels of D-alanylation of teichoic acids and WTA accumulation have been observed in some strains of daptomycin resistant *S. aureus* (more commonly referred to as daptomycin non-susceptible), although their role in antibiotic susceptibility was unclear [40,41]. Based on our findings, we hypothesise that D-alanylated WTA contributes to daptomycin resistance by enabling peptidoglycan accumulation [34]. Whilst this hypothesis would not explain all daptomycin resistance, it is supported by reports describing increased cell wall thickness in conjunction with increased levels of WTA and D-alanylation in some daptomycin non-susceptible strains [47,48,49,50,51].

We confirmed that peptidoglycan accumulation was required for daptomycin tolerance by partial digestion of this polymer with lysostaphin. This finding contrasts with a previous study that showed complete removal of the cell wall led to high levels of daptomycin tolerance in *S. aureus* and *E. faecalis* [28]. A potential reason for these differing findings is that our work only partially removed the wall from growth-arrested bacteria in growth media, whereas the previous study completely removed the wall from growing cells in isotonic buffer (20% sucrose) [28]. Importantly, however, this previous work also showed that inhibition of peptidoglycan synthesis using fosfomycin increased the anti-bacterial activity of daptomycin, in agreement with our work [28].

Whilst there is clearly a key role for the cell wall in daptomycin tolerance caused by growth arrest, inhibition of peptidoglycan did not completely restore susceptibility to the levels seen in replicating cells, indicating that other mechanisms may contribute. For example, membrane fluidity changes as cells enter stationary phase and this alters daptomycin susceptibility [29]. However, this work suggested that daptomycin binding to the membrane was unaltered, which contrasts with our observations here [29]. As such, additional mechanisms by which growth inhibition reduce daptomycin susceptibility remain to be determined.

We have previously found that human serum triggers daptomycin tolerance in *S. aureus* via two independent mechanisms; increased abundance of peptidoglycan and changes to membrane phospholipid composition [34]. Although serum restricts staphylococcal growth, a key difference between daptomycin tolerance induced by growth arrest via antibiotics and that which occurs in serum is the absence of a role for changes to membrane phospholipid composition mediated by Cls2. Whilst cardiolipin was required for serum-induced tolerance [34], we found no evidence that daptomycin tolerance required changes to phospholipid composition. Furthermore, serum-induced tolerance required the presence of the antimicrobial peptide LL-37 and was not triggered simply via inhibition of *S. aureus* growth [34]. Taken together, it appears that daptomycin tolerance can arise via distinct mechanisms in response to various conditions, but in all cases, it appears to require active remodelling of the cell envelope, rather than a lack of metabolic activity.

The reason why some antibiotics kill bacteria and others inhibit growth remains the subject of debate almost 80 years after penicillin was first used clinically [4,5,6,7]. Whilst there is currently no evidence to indicate that bactericidal antibiotics lead to better outcomes for patients than bacteriostatic drugs [52,53,54,55], there is growing evidence that antibiotic tolerance is an underappreciated caused of treatment failure [13,15,56,57,58]. Antibiotic tolerance has also been shown to be a stepping-stone to resistance [16,17]. A recent clinical example of this comes from a patient with relapsing MRSA bacteraemia, which was initially caused by a daptomycin tolerant strain but gave rise to isolates that were daptomycin non-susceptible during subsequent periods of relapse [59].

In addition to the potential clinical implications of antibiotic tolerance, it is anticipated that a better understanding of how antibiotics work and why certain combinations are synergistic may lead to more effective treatments. For example, the inhibition of tolerance by fosfomycin or cephalexin suggests that they may have therapeutic value if used in combination with daptomycin. Indeed, several previous studies have shown synergy between daptomycin and fosfomycin or β-lactams *in vitro*, and our group showed that fosfomycin partially inhibited daptomycin tolerance induced by human serum [34,60,61,62]. There are also promising data from clinical studies, with the presence of a β-lactam or fosfomycin appearing to promote clearance of *S. aureus* infections [63,64,65]. However, combination therapy was generally more nephrotoxic than daptomycin alone and there was very little evidence to suggest enhanced patient survival when daptomycin was used with a cell wall-targeting agent [63,64,65]. Therefore, more work is needed to find the least toxic daptomycin combination therapy to extract clinical benefit [66].

In summary, we have shown that growth arrest results in tolerance to the antibiotic daptomycin, just as occurs for many other bactericidal antibiotics. However, whilst growth arrest typically confers antibiotic tolerance passively via reduced metabolic activity, daptomycin tolerance arises via the active synthesis and accumulation of peptidoglycan.

## Methods

### Bacterial strains and growth conditions

The bacterial strains used in this study are shown in Table 1. Strains were routinely grown in tryptic soy broth (TSB; BD Biosciences) for 16 h at 37 ° C with shaking (180 rpm) or on tryptic soy agar (TSA; BD Biosciences). Where appropriate, media were supplemented with 10 μg ml^-1^ erythromycin.

**Table 1.** Strains used in this study.

| Strain | Characteristics | Reference |
| --- | --- | --- |
| USA300 LAC* | LAC strain of the USA300 CA-MRSA lineage, cured of LAC-p03 plasmid | 67 |
| USA300 LAC* $\Delta tagO$ | USA300 LAC* with the *tagO* gene deleted | 68 |
| USA300 LAC* $\Delta ltaS$-S2 | USA300 LAC* $\Delta ltaS::erm$ suppressor S2, Ery<sup>r</sup> | 69 |
| USA300 LAC* $\Delta dltD$ | USA300 LAC* with the *dltD* gene deleted, Ery<sup>r</sup> | 34 |
| USA300 LAC JE2 | LAC strain of the USA300 CA-MRSA lineage cured of plasmids | 70 |
| USA300 LAC JE2 *mprF*::Tn | USA300 LAC JE2 with a *bursa aurealis* transposon insertion in *mprF*, Ery<sup>r</sup> | 70 |
| USA300 LAC JE2 *cls1*::Tn | USA300 LAC JE2 with a *bursa aurealis* transposon insertion in *cls1*, Ery<sup>r</sup> | 70 |
| USA300 LAC JE2 *cls2*::Tn | USA300 LAC JE2 with a *bursa aurealis* transposon insertion in *cls2*, Ery<sup>r</sup> | 70 |
| USA300 LAC JE2 *crtM*::Tn | USA300 LAC JE2 with a *bursa aurealis* transposon insertion in *crtM*, Ery<sup>r</sup> | 70 |

Ery, erythromycin.

### Generation of mid-exponential phase and growth-arrested cultures

Mid-exponential phase cultures were generated by dilution of overnight cultures to 10^7^ CFU ml^-1^, and then growth for 2 h at 37 ° C with shaking (180 rpm) until 10^8^ CFU ml^-1^ was reached. To generate growth-arrested cultures, bacteriostatic concentrations of the appropriate antibiotic was added (Table 2). Due to different susceptibilities of some mutants to some antibiotics, the concentrations used were optimised for each mutant. Except where stated, growth arrest was performed in TSB for 16 h at 37 *°* C with shaking (180 rpm). As bacteriostatic concentrations were used, the CFU ml^-1^ remained constant (10^8^ CFU ml^-1^). Where appropriate, growth arrest was performed in phosphate buffered saline (PBS), cation-adjusted Müller Hinton broth (MHB) or MHB supplemented with 2.5 g L^-1^ glucose.

**Table 2.** Concentrations of antibiotics used in this study.

| Antibiotic | Supplier | Strain | Concentration ($\mu\text{g ml}^{-1}$) |
| --- | --- | --- | --- |
| Tetracycline (TSB) | Sigma | LAC* WT | 1.25 |
| | | LAC* $\Delta tagO$ | 0.156 |
| | | LAC* $\Delta ltaS$-S2 | 0.078 |
| | | LAC* $\Delta dltD$ | 0.3125 |
|  |  | JE2 WT | 0.625 |
|  |  | JE2 *mprF*::Tn | 0.625 |
|  |  | JE2 *cls1*::Tn | 0.625 |
|  |  | JE2 *cls2*::Tn | 0.625 |
|  |  | JE2 *crtM*::Tn | 0.625 |
| Tetracycline (MHB) | Sigma | LAC* WT | 5 |
| Chloramphenicol | Sigma | LAC* WT | 20 |
| Ciprofloxacin | Sigma | LAC* WT | 160 |
| Cephalexin | Tokyo Chemical Industry | LAC* WT | 32 |
| Tunicamycin | Abcam | LAC* WT | 0.5 |
| AFN-1252 | MedChemExpress | LAC* WT | 0.15 |
| Fosfomycin | Tokyo Chemical Industry | LAC* WT | 8 |
| Amsacrine | Sigma | LAC* WT | 10 |
| Lysostaphin | Sigma | LAC* WT | 1 |

TSB, tryptic soy broth; MHB, Müller Hinton broth.

### Daptomycin killing assays

Mid-exponential phase and growth-arrested cultures (3 ml) were generated as described above. CaCl_2_ was added to a final concentration of 1.25 mM and daptomycin was added to 20 μg ml^-1^. Cultures were incubated at 37 ° C with shaking (180 rpm) for 6 h. At each time point, aliquots were taken, serially diluted 10-fold in PBS and plated onto TSB to determine CFU ml^-1^. Where appropriate, after growth arrest cells were incubated with 1 μg ml^-1^ lysostaphin in TSB for 1 h at 37 ° C to partially remove the cell wall before addition of daptomycin.

### Measurements of daptomycin binding

Daptomycin was labelled with the BoDipy fluorophore as described previously [34]. Cultures (3 ml) of exponential phase or growth-arrested bacteria were incubated at 37 ° C with shaking (180 rpm) with 80 μg ml^-1^ BoDipy-daptomycin in TSB supplemented with 1.25 mM CaCl_2_. Every 2 h, aliquots were taken and washed in PBS three times. Samples (200 μl) were put into black-walled flat-bottomed 96 well plates and fluorescence measured using a TECAN Infinite 200 PRO microplate reader (excitation 490 nm; emission 525 nm). Alternatively, daptomycin binding was investigated by fluorescence microscopy as described below.

### Measurements of membrane depolarisation

Membrane polarity was measuring using 3,3’-dipropylthiadicarbocyanine iodide (DiSC3(5); Thermofisher Scientific), as described previously [34]. This dye binds to polarised membranes, quenching its fluorescence and resulting in higher fluorescence values correlating with increased membrane depolarisation. Exponential phase and growth-arrested cultures were exposed to 20 μg ml^-1^ daptomycin for 6 h at 37 ° C in 3 ml TSB supplemented with 1.25 mM CaCl_2_. Every 2 h, 200 μl aliquots were taken into black-walled, flat-bottomed 96-well plates. DiSC3(5) was added to a final concentration of 1 μM, mixed and the plate incubated statically at 37 ° C for 5 min. Fluorescence was measured using a TECAN Infinite 200 PRO microplate reader (excitation 622 nm; emission 670 nm) and values were divided by OD_600_ measurements to normalise for differences in cell density.

### Measurements of membrane permeability

Membrane permeability was measured using propidium iodide (PI; Sigma), a membrane-impermeant dye which fluoresces when bound to DNA, as described previously [34]. Exponential phase and growth-arrested cultures were exposed to 20 μg ml^-1^ daptomycin for 6 h at 37 ° C in 3 ml TSB supplemented with 1.25 mM CaCl_2_. Every 2 h, 200 μl aliquots were washed three times with PBS, added into a black-walled, flat-bottomed 96-well plate and PI added to a final concentration of 2.5 μM. Fluorescence was measured using a TECAN Infinite 200 PRO microplate reader (excitation 535 nm; emission 617 nm) and values were divided by OD_600_ measurements to normalise for differences in cell density.

### Phase contrast and fluorescence microscopy

Exponential phase or growth-arrested cultures were incubated with BoDipy-daptomycin as described above. After 2 h, aliquots were taken, washed three times in PBS and fixed in 4 % paraformaldehyde. To measure peptidoglycan synthesis, exponential phase and growth-arrested cultures were generated with the addition of 25 μM HADA. Samples were washed three times in PBS and fixed in 4 % paraformaldehyde.

Aliquots of fixed bacteria (2 μl) were spotted onto agarose (1.2 % in water) on microscope slides and covered with a cover slip. Images were taken with a Zeiss Axio Imager.A1 microscope coupled to an AxioCam MRm and a 100x objective. BoDipy-daptomycin was detected using a green fluorescent protein filter set and HADA using a DAPI filter set. Images were processed using the Zen 2012 software (blue edition). Within an experiment, microscopy of all samples was performed at the same time using identical settings to enable comparisons to be made between samples.

### Extraction and quantification of WTA

WTA was extracted from 40 ml cultures of exponential phase or growth-arrested cultures as described previously [34]. Briefly, cultures were washed with 50 mM MES, pH 6.5, and incubated at 100 ° C for 1 h in 4 % SDS, 50 mM MES, pH 6.5. Samples were then washed twice in 4 % SDS, 50 mM MES, once with 2 % NaCl, 50 mM MES and once with 50 mM MES before being resuspended in 1 ml 20 mM Tris-HCl pH 8, 0.5 % SDS, 20 μg ml^-1^ proteinase K and incubated for 4 h at 50 ° C (shaking at 1400 rpm). The pellet was recovered, washed with 2 % NaCl, 50 mM MES and three times with water before being resuspended in 1 ml 0.1 M NaOH and incubated at 20 °C for 16 h (shaking at 1400 rpm). After centrifugation at 16,000 x *g* for 1 min, the supernatant was neutralised with 250 μl 1 M Tris-HCl, pH 7.8 and stored at −20 ° C.

Purified WTA was loaded on 20 % native polyacrylamide gels and run at 120 V in Tris-Tricine running buffer (0.1 M Tris, 0.1 M Tricine) before being stained with 0.1 % alcian blue in 3 % acetic acid. Gels were destained with water and imaged using a Gel Doc EZ Imager (Bio-Rad). WTA intensity was quantified using ImageJ.

### Extraction, detection and quantification of LTA

Exponential phase and growth-arrested cultures (10 ml) were resuspended in 1 ml PBS and transferred to screw cap tube containing approximately 100 μl 0.1 mm glass beads. Cells were lysed at room temperature using a FastPrep-24 (MP Biomedicals) machine (4 cycles of 6.5 m/s for 40 s followed by a 1 min rest). Tubes were centrifuged at 200 x *g* for 1 min to settle the beads and 500 μl supernatant removed into a new tube. Centrifugation (16,000 x *g* for 15 min) pelleted the cellular debris including the LTA. The supernatant was discarded and the pellet resuspended in 100 μl 2x Laemmli sample buffer (4 % SDS, 20 % glycerol, 10 % β-mercaptoethanol, 0.02 % bromophenol blue, 0.125 M Tris-HCl, pH 6.8). Samples were incubated at 95 ° C for 20 min, then centrifuged at 17,000 x *g* for 5 min before the supernatant, containing LTA, was moved to a fresh Eppendorf and stored at −20 C.

LTA extracts (10 μl) were run on 15 % polyacrylamide gels and transferred to PVDF membranes using a Trans-Blot Turbo transfer system (Bio-Rad). After blocking with 5 % milk and 1 % BSA in TBST, LTA was detected with an anti-LTA primary antibody (mAb 55; HycultBiotech; 1:5000 dilution) and an HRP-goat anti-mouse IgG secondary antibody (Abcam; 1:10000 dilution). LTA was visualised using Amersham ECL Prime western blotting detection reagent (GE Healthcare) and a ChemiDoc imaging system (Bio-Rad). LTA intensity was quantified using ImageJ.

### Measurement of surface charge

Bacterial surface charge was determined using fluorescein isothiocyanate-labelled poly-L-lysine (FITC-PLL). Aliquots (1 ml) of exponential phase and growth-arrested cultures were incubated with 80 μg ml^-1^ FITC-PLL for 10 min at room temperature in the dark before being washed three times in PBS. 200 μl was then moved to a black-walled 96 well plate and fluorescence measured using a TECAN Infinite PRO 200 plate reader (excitation 485 nm; emission 525 nm).

### Measurements of peptidoglycan incorporation by plate reader

Peptidoglycan synthesis was measured using HCC-amino-D-alanine (HADA), a fluorescent D-amino acid analogue. Exponential phase and growth-arrested cultures were generated with the addition of 25 μM HADA. Samples were washed three times in PBS and 200 μl aliquots moved to black-walled 96 well plates. Fluorescence was measured using a TECAN Infinite 200 PRO microplate reader (excitation 405 nm; emission 450 nm).

### Statistical analyses

CFU data were log_10_ transformed. Data were analysed by Student’s *t*-test, one-way ANOVA or two-way ANOVA with a *post-hoc* test to correct for multiple comparisons as described in the figure legends using GraphPad Prism (v8.0).

## Acknowledgements

Angelika Grundling (Imperial College London) thanked for providing strains. Simon Foster (University of Sheffield) is thanked for providing HADA. Vladimir Pelicic (Imperial College London) is thanked for providing access to microscopy equipment. All authors acknowledge the provision of strains by the Network on Antimicrobial Resistance in *Staphylococcus aureus* (NARSA) Program: under NIAID/ NIH Contract No. HHSN272200700055C. The funders had no role in the study design, interpretation of the findings or the writing of the manuscript.

## Contributions

EVKL conducted experiments. AME and EVKL designed experiments, analysed data and wrote the manuscript.

